# Microorganisms consume trace gases present throughout the Mariana Trench water column

**DOI:** 10.64898/2026.09.23.753970

**Authors:** Yuka Katayama, Francesco Ricci, Yuto Fukuyama, Eiji Tasumi, Yukari Yoshida-Takashima, Akiko Makabe, Takuro Nunoura, Taichi Yokokawa, Ken Takai, Chris Greening

## Abstract

Dark waters harbour two-thirds of the ocean’s bacterial and archaeal cells, yet the energy sources sustaining microbial life at these depths remain poorly understood. Here we integrate hydrographic profiles with metagenomic analyses and *ex situ* activity measurements to investigate whether carbon monoxide (CO) and molecular hydrogen (H_2_) are viable energy sources for epipelagic and mesopelagic microbial populations in the Mariana Trench region. Both gases were detected from 5 to 9,371 m, with concentrations generally higher in the photic zone but anomalously high in several deeper waters. Metagenomic analyses indicate a shift from phototrophic towards lithotrophic energy acquisition with depth. Microbial populations encoding CO dehydrogenases increased from 6.1% at 5 m to 45% at 1,000 m, while those encoding uptake hydrogenases were much less abundant, but increased from 0.34% to 0.63%. *Ex situ* incubations revealed that *in situ* cell-specific CO oxidation rates declined with depth, whereas corresponding H_2_ oxidation rates were significantly higher at 1,000 m than at the surface. At 1,000 m, cell-specific H_2_ oxidation rates were approximately 4,500-fold higher than those of CO. These contrasting trends suggest that CO oxidation is a widespread auxiliary energy-acquisition strategy, whereas H_2_ oxidation may sustain hydrogenotrophic growth in dark waters. Together, our results suggest H_2_ and CO are readily available to pelagic microbial populations and are biologically oxidised in the dark zone, supporting microbial energy acquisition and oceanic biogeochemical cycling.

## Main

The dark ocean, comprising water below 200 m, is the largest habitat in the biosphere by volume and harbours nearly 70% of oceanic bacterial and archaeal biomass in ocean waters [1]. As light and organic carbon supply decline with depth, chemolithotrophy becomes increasingly important [2], yet the inorganic energy sources supporting microbial life in the dark water column remain incompletely understood. Carbon monoxide (CO) and molecular hydrogen (H_2_) are ubiquitous and energy-rich trace gases that support microbial growth and long-term survival [3]. Although the ocean is a net atmospheric sources of both gases, microbial oxidation constitutes a major internal sink, accounting for ∼85–99% of CO [4, 5] and

∼99% of H_2_ removal in studied marine regions [6]. Their oxidation is mainly mediated by oxygen-tolerant molybdenum-dependent CO dehydrogenase (Mo-CODH) and [NiFe]-hydrogenases, respectively, which are encoded by phylogenetically and metabolically diverse microorganisms [3]. Across global ocean metagenomes, genes encoding these enzymes occur in an estimated average of 10.4% and 0.8% of microbial genomes and become more prevalent with depth [7]. Despite this widespread genetic potential, evidence for trace gas utilization in the deep ocean remains sparse. Although background-water CO and H_2_ concentrations have been measured to 4,500 m [8] and 3,230 m [9], respectively, such profiles remain limited, with none reported from the Mariana Trench. More importantly, biological CO and H_2_ oxidation in background seawater have been directly demonstrated to 120 m [10] and 25 m [11] to our knowledge, respectively, leaving trace gas oxidation in the dark water column undetected.

Here, we investigated CO and H_2_ availability, and microbial functional potential and oxidation activity in epipelagic and mesopelagic waters overlying the Challenger and Sirena Deeps of the Mariana Trench. These are the first and third deepest parts of the ocean in the world and were surveyed during the R/V *Kaimei* cruise KM25-12 in 2026 (JAMSTEC) [12] (Fig. 1A; Table S1; Supplementary Methods). Oxygen and nitrate were detected in pelagic waters, as reported previously in the region [13] (Fig. 1B). We quantified the levels of H_2_ and CO in the water column from 5 to 9,371 m through highly sensitive gas chromatography. Both gases were detected across all depths (Fig. 1C,D; Table S2). CO concentrations were highest at 5– 6 m (0.53–0.74 nM), reaching up to 7.3-fold air-sea equilibrium, but declined to 0.07–0.15 nM by 150 m and remained mostly between 0.04 and 0.11 nM below 200 m (Fig. 1C). This gradient is consistent with photochemical production near the surface followed by biological removal [5]. The presence of CO throughout dark and hadal waters nevertheless indicates continued supply through transport and/or dark production. The Challenger Deep additionally showed a distinct enrichment between 6,500 and 7,500 m, with CO peaking at 0.23 nM at 7,000 m in contrast to the Sirena Deep, likely reflecting the impacts of subseafloor fluids on the trench system. H_2_ concentrations were higher and more variable than CO, ranging from 0.17 to 4.0 nM (Fig. 1D). Both H_2_ full-depth profiles showed a common structure, with supersaturation in the photic zone and declining toward seafloor with several anomalies. The surface levels are likely due to H_2_ production during cyanobacterial nitrogen fixation and dissolved organic matter photochemistry, with nitrogen fixation and fermentation in dark microenvironments [14, 15], potentially driving supersaturation of this gas in other depths.

**Fig. 1.**
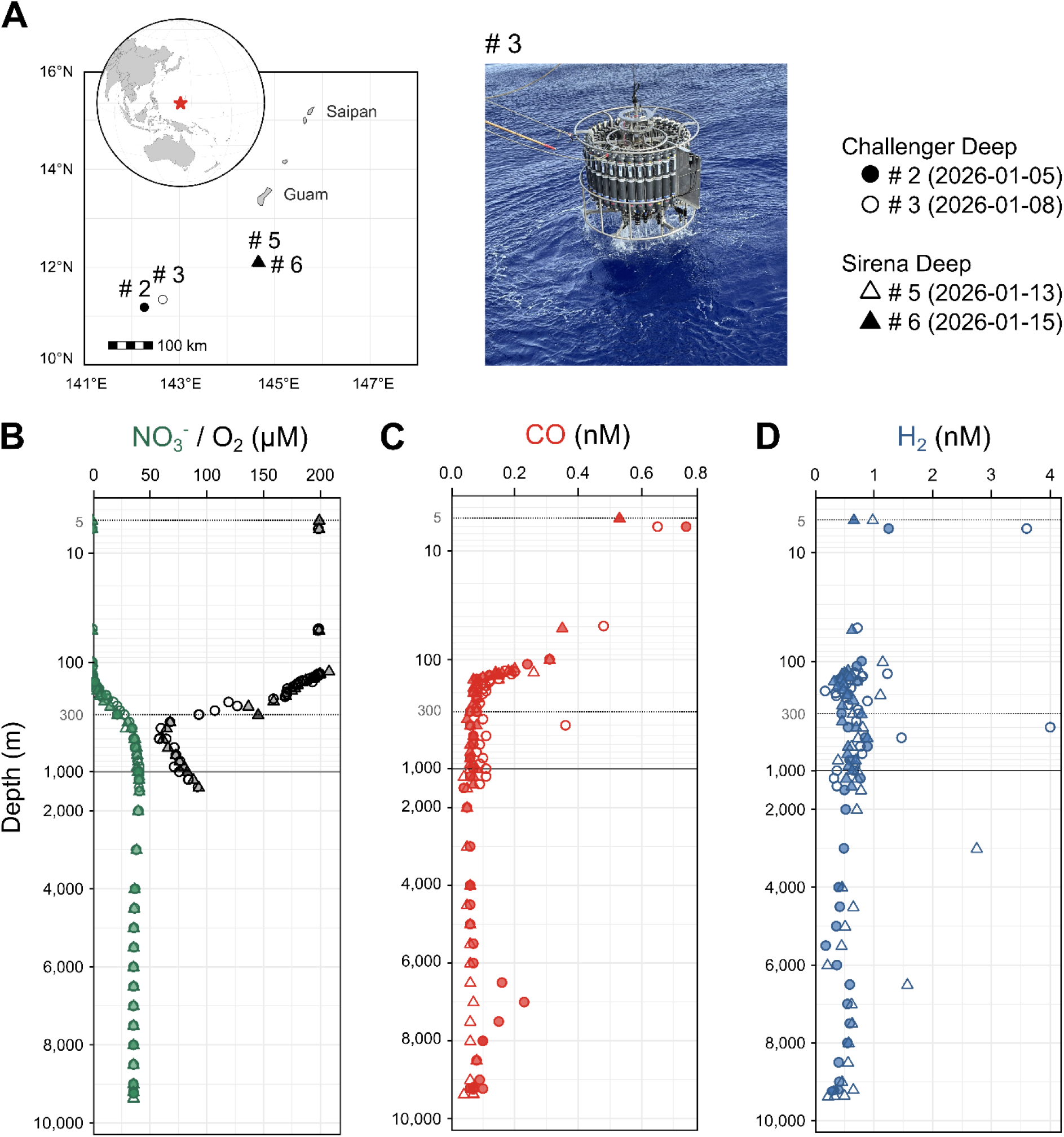
Water-column distributions of carbon monoxide, molecular hydrogen, and electron acceptors in the Mariana Trench. (A) Locations of sampling stations overlying the Challenger Deep (stations #2 and #3, circles) and Sirena Deep (stations #5 and #6, triangles) in the western North Pacific Ocean. Filled and open symbols distinguish individual casts as indicated. The inset shows the location of the Mariana Trench, and the photograph shows CTD-rosette deployment at station #3. (B) Depth profiles of nitrate (green) and dissolved oxygen measured to 1,400 m (black). (C, D) Depth profiles of dissolved carbon monoxide (CO) and molecular hydrogen (H_2_) measured from 5 m to maximum depths of 9,241 m in Challenger Deep and 9,371 m in Sirena Deep. Symbols correspond to the stations shown in (A). Each point represents an individual seawater measurement. Depth axes use a logarithmic scale above 1,000 m and a linear scale below 1,000 m. Horizontal dotted lines at 5 and 300 m and the solid line at 1,000 m mark the nominal sampling depths for metagenomic analyses and activity assays.

We used metagenomics to profile the capabilities of the microbial communities inhabiting these waters, focusing on the upper 1,000 m water column given limited shipboard experimental capacity. Short-read metagenomes from 5 m (epipelagic), 300 m (upper mesopelagic), and 1,000 m (lower mesopelagic) waters revealed concordant community structure and shifts at both the Sirena and the Challenger Deep. Epipelagic communities harboured sequences associated with Cyanobacteriota (Fig. 2A) and enrichment of genes involved in light-driven energy conservation, the Calvin-Benson cycle, and nitrogen fixation (Fig. S1; Table S3). In mesopelagic waters, these features declined as sequences associated with lineages such as Thermoproteota and Chloroflexota increased alongside light-independent energy metabolisms. This indicates a transition from phototrophic production towards increasing reliance on organic matter remineralisation and inorganic energy acquisition. This transition was accompanied by an increase in relative abundance of trace gas oxidation genes (Fig. 2B,C).

**Fig. 2.**
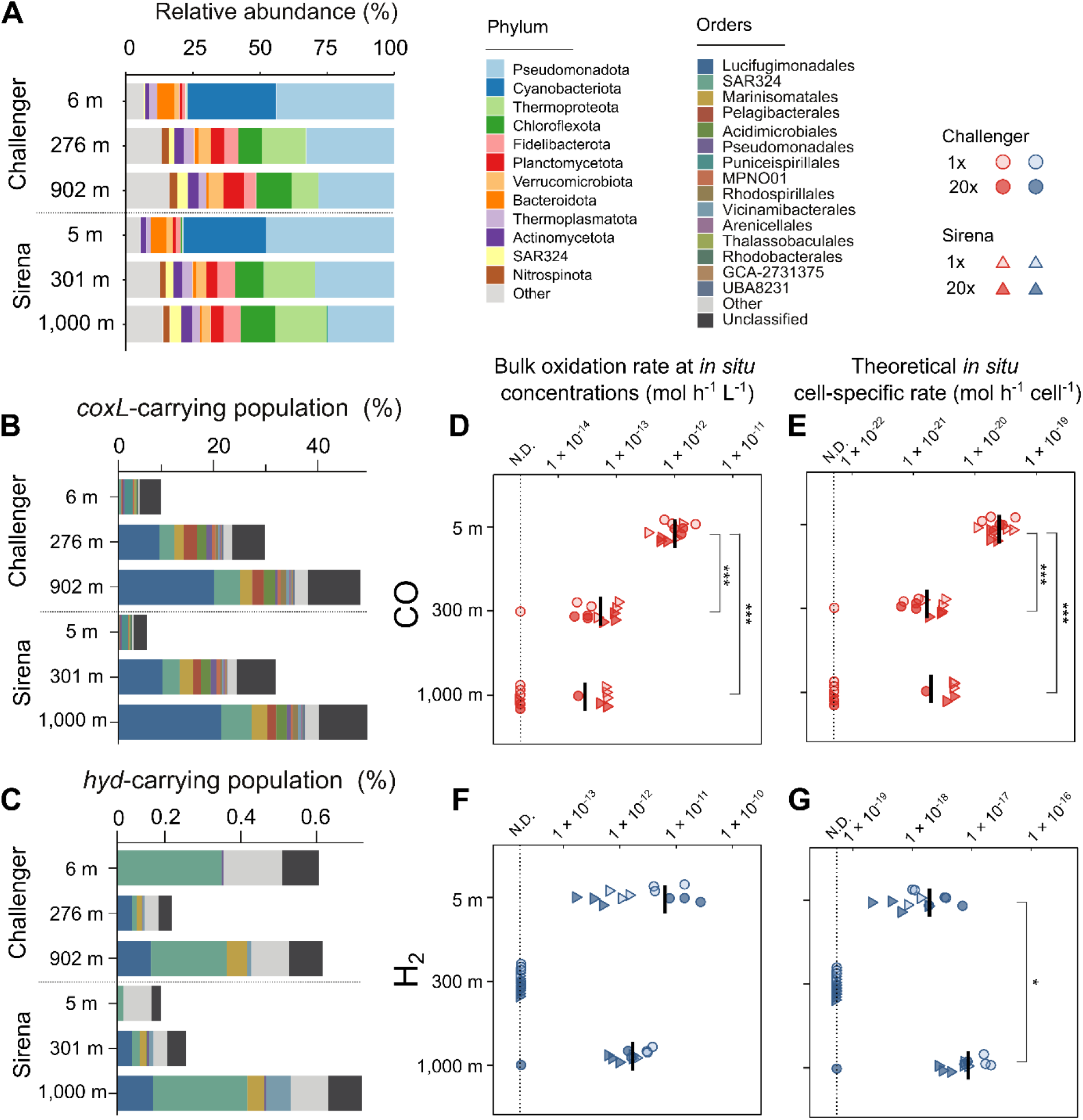
Depth-dependent microbial communities, trace gas oxidation potential, and activity. (A) Phylum-level composition of prokaryotic communities reconstructed from short read metagenomes collected at 5-6 m, approximately 300 m, and approximately 1,000 m overlying the Challenger and Sirena Deeps. (B, C) Estimated prevalence and order-level composition of populations carrying form I Mo-CODH (*coxL*) (B) or group 1 and 2 [NiFe]-hydrogenases (*hyd*) (C). Bar lengths indicate the estimated proportion of the community carrying the corresponding genes, and colours indicate taxonomic orders. (D, E) CO oxidation and (F, G) H_2_ oxidation. Panels show bulk oxidation rates scaled to measured *in situ* gas concentrations (D, F) and theoretical *in situ* cell-specific rates normalized to the estimated abundance of cells carrying *coxL* or *cooS* (E) or group 1 and 2 [NiFe]-hydrogenase genes (G). Circles and triangles indicate Challenger and Sirena Deep samples, respectively; open and filled symbols indicate incubations with native seawater and cell concentrates prepared from a 20-fold larger seawater volume, respectively. Rates from concentrated-cell incubations were corrected using the measured cell concentration factors to express activity on a native-seawater basis. Each symbol represents an individual microcosm, and horizontal black bars indicate means. N.D. denotes incubations in which gas consumption was not detected relative to the paired heat-killed control; these values are displayed on the horizontal dotted line for visualization and were treated as zero in statistical analyses. Differences among depths were evaluated using the Kruskal-Wallis test followed by Dunn’s post hoc test with Holm-Bonferroni adjustment. \**P* < 0.05; \*\*\**P* < 0.0001.

The estimated mean prevalence of microbes encoding form I Mo-CODH (*coxL*) increased from 6.1% at 5 m to 27% at ∼300 m and 45% at ∼1,000 m (Fig. 2B). Surface *coxL*-containing taxa were predominantly aerobic heterotrophs, including Pseudomonadales (Gammaproteobacteria) and Puniceispirillales and Thalassobaculales (Alphaproteobacteria). Deeper *coxL*-containing taxa shifted towards oligotrophic and metabolically versatile Lucifugimonadales (Chloroflexota), SAR324, Acidimicrobiales (Actinomycetota) and Marinisomatales (Marinisomatota), associated with organic matter remineralisation and chemolithotrophy [16, 17] (Fig 2B; Table S4), suggesting that CO oxidation provides energy to support diverse metabolic strategies in dark zones. In contrast, oxygen-sensitive nickel-dependent CODH (*cooS*) was rare and detected only at 1,000 m in both Deeps (Table S3). Microbes encoding group 1 and 2 [NiFe]-hydrogenases were much less abundant, with mean estimated prevalence of 0.34% at 5 m, 0.17% at 300 m and 0.63% at 1,000 m (Fig. 2C; Table S3). High-affinity group 1l [NiFe]-hydrogenases dominated throughout, while surface-specific group 2a [NiFe]-hydrogenases and abundant Cyanobacteriota suggest nitrogen fixation-associated H_2_ recycling [18]. Deeper waters additionally contained group 1d [NiFe]-hydrogenases associated with aerobic hydrogenotrophic growth [3], alongside groups 1b and 1h. Putative H_2_ oxidisers were represented by SAR324 throughout the water column (Fig 2C). Surface waters also harboured Phycisphaerae (Planctomycetota) and Actinomarinales (Actinomycetota), whereas deeper putative H_2_ oxidizers include Lucifugimonadales (Chloroflexota), Marinisomatales (Marinisomatota), Vicinamibacterales (Acidobacteriota), and UBA8108 (Planctomycetota) (Fig 2C; Table S4). Together, these patterns suggest a shift from nitrogen fixation-associated H_2_ recycling at the surface towards phylogenetically diverse persistence- and growth-linked H_2_ oxidation in dark waters.

We next tested whether trace gas oxidation functional potential translated into measurable activity (Fig. 2D,E; Fig. S2; Table S5). Native seawater and cell concentrates prepared from a 20-fold larger volume of seawater were incubated with headspaces amended to 2 ppm CO and H_2_ at near *in situ* temperatures (See Supplementary Methods). CO oxidation was detected in 12/12, 11/12, and 6/12 incubations at 5, 300, and 1,000 m, respectively. Scaled *in situ* mean bulk rates showed a significant decline with depth from 1.0 × 10^-12^ mol L^-1^ h^-1^ at 5 m to 5.4 × 10^-14^ and 2.9 × 10^-14^ mol L^-1^ h^-1^ at 300 and 1,000 m (Fig. 2D). Mean theoretical *in situ* cell-specific CO oxidation rates per *coxL/cooS*-carrying cell were 15-fold lower at 300 m and 13-fold lower at 1,000 m than at 5 m (Fig. 2E). H_2_ oxidation showed a contrasting pattern, being detected in 12/12 incubations at 5 m, none at 300 m, and 11/12 at 1,000 m. Mean *in situ* bulk rates were 6.3 × 10^-12^ and 1.71 × 10^-12^ mol L^-1^ h^-1^ at 5 and 1,000 m, respectively, with no significant difference between depths (Fig. 2F). Notably, mean theoretical *in situ* cell-specific H_2_ oxidation rates were 80-fold and 4,523-fold higher than those of CO at 5 and 1,000 m, respectively (Fig. 2E,G; Table S5). Post-incubation enrichment of *Labrenzia* at 5 and 300 m and *Sulfurimonas* at 1,000 m, genera containing experimentally verified CO- and H_2_-oxidizing members, respectively [19, 20], may partly account for the observed activity patterns (Table S6). The absence of detectable H_2_ oxidation in the upper mesopelagic was unexpected, but consistent with the minimal levels of hydrogenase genes at this depth, suggesting complex environmental controls influence the distribution and activity of trace gas oxidizers.

Together, our findings show that CO and H_2_ are biologically oxidized in ultra-deep trench waters to at least 1,000 m and are present as energy sources throughout the 9,371-m Mariana water column. H_2_ oxidation may make a small but substantial contribution to the energy budget of dark waters, with elevated cell-specific rates at the bottom of mesopelagic zone, whereas the contribution of CO oxidation is limited in dark water despite the widespread distribution of CO dehydrogenase genes in mesopelagic microbial communities. CO oxidation rates were lower than reported elsewhere [5, 7, 10], potentially reflecting the oligotrophic setting, whereas H_2_ oxidation remained comparable between surface and 1,000-m waters. These observations extend the surface measurements and depth-resolved genomic predictions of Lappan *et al*. (2023) with the first direct demonstration of CO and H_2_ oxidation in dark oceanic water columns to our knowledge. The ubiquity of trace gases in deep waters suggests trace gas oxidation is a previously overlooked dimension of microbial energy acquisition in Earth’s largest habitat.

## Data availability

The raw reads for metagenome were deposited in the NCBI Sequence Read Archive under Bioproject accession number PRJNA1530696.

## Funding

This work was supported by an ARC Discovery Project grant (DP260101905 to C.G. and F.R.) and an ARC Future Fellowship (FT240100502 to C.G.). Y.K. is supported by JSPS Overseas Research Fellowship.

## Cruise Permission

We declare that KM25-12 cruise was conducted under the special conditions and requirements of the United States Department of State, and of the Marianas Trench National Wildlife Refuge and Mariana Arc of Fire National Wildlife Refuge of the U.S. Fish and Wildlife Service with a permission SUP#12540-25002 and under the special conditions and requirements of the Government of the Federated States of Micronesia with a permission FM25-JP25001RS-26266.

## Acknowledgements

We thank Masayuki Miyazaki for assistance with the incubation experiments and Dr. Tess Hutchinson for technical advice. We also thank the captain and crew of R/V *Kaimei* and the staff of JAMSTEC and Marine Works Japan Ltd. for their support during cruise KM25-12.

## Author contributions

Y.K. and C.G. conceived and designed the study. T.N. facilitated the collaboration and contributed to experimental planning. K.T. led the KM25-12 research cruise. Y.F. coordinated project implementation and provided extensive technical and logistical support, while T.Y. led the seawater research component and sampling operations. Y.K., Y.F., E.T., A.M. and T.Y. performed field sampling. Y.K. conducted the incubation experiments and performed trace gas oxidation and amplicon analyses. E.T. conducted dissolved-gas measurements, A.M. conducted nutrient measurements, Y.Y. performed metagenomic and amplicon sequencing, and F.R. performed metagenomic analyses. Y.K. and F.R. drafted the manuscript, and C.G. contributed to manuscript revision. All authors reviewed and approved the final manuscript.

## Conflict of interest

The authors declare that they have no conflict of interest.

## Notes

https://www.ncbi.nlm.nih.gov/bioproject?term=PRJNA1530696

### Competing Interest Statement

The authors have declared no competing interest.

