## Supplementary material for "Microorganisms consume trace gases present throughout the Mariana Trench water column": Fig. S

### **Supplementary Information for** **Microorganisms consume trace gases** **present throughout the Mariana Trench water column**

Yuka Katayama<sup>1</sup>, Francesco Ricci<sup>1</sup>, Yuto Fukuyama<sup>2</sup>, Eiji Tasumi<sup>2</sup>, Yukari Yoshida-Takashima<sup>2</sup>,
Akiko Makabe<sup>2</sup>, Takuro Nunoura<sup>2</sup>, Taichi Yokokawa<sup>2,3</sup>, Ken Takai<sup>2,\*</sup>, Chris Greening<sup>1,\*</sup>

<sup>1</sup>Department of Microbiology, Biomedicine Discovery Institute, Monash University, Clayton, VIC 3800, Australia

<sup>2</sup>Institute for Extra-cutting-edge Science and Technology Avant-garde Research of Life (X-star), Japan Agency for Marine-Earth Science and Technology (JAMSTEC), 2–15 Natsushima-cho, Yokosuka, Kanagawa, 237–0061, Japan

<sup>3</sup>Advanced Institute for Marine Ecosystem Change (WPI-AIMEC), JAMSTEC, 2–15 Natsushima-cho, Yokosuka, Kanagawa, 237–0061, Japan

#### **Materials and Methods**

##### **Sample collection**

The seawater samples used in this study were obtained by vertical CTD hydrocasts during the R/V *Kaimei* KM25-12 cruise in January 2026 (JAMSTEC; [1]), and the sampling stations and water depths are summarized in Table S1 and Fig. 1A. Hydrocasts at stations #2 and #3 in Challenger Deep were conducted on 5 and 8 January, respectively, and those at stations #5 and #6 in Sirena Deep were conducted on 13 and 15 January, respectively. The full-depth hydrocasts at stations #2 and #6 reached maximum depths of 9,241 and 9,371 m, respectively, whereas the hydrocasts at stations #3 and #5 covered the upper water column to maximum depths of 1,413 and 1,500 m, respectively.

The CTD frame used for all hydrocasts included an SBE 9plus pressure sensor, an SBE 3Plus temperature sensor, and an SBE 4C conductivity sensor (Sea-Bird Scientific, USA). The upper-water hydrocasts additionally included an SBE 43 dissolved oxygen sensor and a C-Star transmissometer (Sea-Bird Scientific, USA), and a chlorophyll fluorometer (Seapoint Sensors Inc., USA). Sensors rated to a maximum depth of 6,000 m were removed from the CTD frame during the full-depth hydrocasts. Seawater samples were collected for shore-based chemical and microbiological analyses, including CO and H<sub>2</sub> measurements, nutrient analyses, microbial cell counting, and metagenomic sequencing.

##### **Dissolved carbon monoxide and hydrogen measurements**

Dissolved CO and H<sub>2</sub> were sampled and analysed as previously reported [2], with minor modifications. Seawater was transferred directly from the Niskin bottles into duplicate 120-mL glass vials through PTFE tubing, avoiding bubble formation and allowing more than two vial volumes to overflow. Each vial was amended with 0.5 mL of saturated HgCl<sub>2</sub> solution and sealed with a PTFE-lined butyl-rubber septum and an aluminum crimp. The samples were stored in the dark at approximately 5°C until analysis. A 20-mL headspace was created by replacing an equal volume of seawater with nitrogen using gas-tight syringes. The vials were shaken for 6 min at 22°C to equilibrate the dissolved gases with the headspace. A 1-mL aliquot of the equilibrated headspace was analyzed using a GC-8A gas chromatograph (Shimadzu, Kyoto, Japan) equipped with a trace reduced-gas detector (TRD-1; Round Science Inc., Japan). CO and H<sub>2</sub> were separated using a 3-m Unibeads C

column (mesh 60/80, I.D. 2.2mm) and a 3-m Molecular Sieve 13X-S column (mesh 80/100, I.D. 2.2mm) maintained at 40°C with nitrogen as a carrier gas. Instrument responses were calibrated against a standard gas mixture containing (i) 1.06 ppm CO and 1.00 ppm H<sub>2</sub> and (ii) 10.4 ppm CO and 10.1 ppm H<sub>2</sub> (Taiyonsan, Tokyo, Japan). Dissolved concentrations were calculated from the measured headspace mixing ratios by mass balance using the Bunsen solubility coefficients for seawater at a salinity of 34 ppt and an equilibration temperature of 22°C [3].

##### **Nitrate and other nutrient measurements**

Seawater samples for nutrient analysis were collected in plastic tubes and stored at –20°C until analysis. Concentrations of nitrate, nitrite, phosphate, and silicate were determined spectrophotometrically using an automated analyzer (QuAAtro39-J; BL TEC, Japan), according to established methods [4–7].

##### **Microbial cell counting**

Samples were fixed with glutaraldehyde at a final concentration of 0.5%, transferred to 2-mL cryovials (Nalgene), and stored at –80°C until analysis [8, 9]. The abundances of prokaryotes, virus-like particles (VLPs), and phytoplankton were determined using an Attune NxT flow cytometer (Thermo Fisher Scientific). For prokaryotic enumeration, a 200-μL aliquot of each thawed sample was stained with SYBR Green I (Invitrogen) at a final dilution of $5 \times 10^{-4}$ of the commercial stock solution and incubated for at least 10 min at room temperature in the dark. A 75-μL aliquot of the stained sample was analyzed using an Attune NxT flow cytometer (Thermo Fisher Scientific) with excitation at 488 nm and fluorescence detection through 530/30-nm (BL1) and 574/26-nm (BL2) bandpass filters. Prokaryotic cells were identified based on their fluorescence and side-scatter characteristics, following Yang et al. (2010). For VLPs enumeration, thawed samples were diluted 20- to 100-fold in TE buffer (10 mmol L<sup>-1</sup> Tris-HCl and 1 mmol L<sup>-1</sup> ethylenediaminetetraacetic acid, pH 8.0; Wako). The diluted samples were stained with SYBR Green I at a final dilution of $5 \times 10^{-5}$ of the commercial stock solution and incubated at 80°C for 10 min. VLPs were identified and enumerated based on their fluorescence and side-scatter characteristics as described [9]. For phytoplankton enumeration, thawed samples were analyzed without staining. *Prochlorococcus*, *Synechococcus*, and photosynthetic picoeukaryotes were distinguished based on their autofluorescence and light-scattering characteristics using cytograms of red fluorescence (BL3: 695/40-nm bandpass filter) versus side scatter, orange fluorescence (BL2) versus side scatter, orange versus red fluorescence, and side scatter versus forward scatter [10].

##### **DNA extraction and metagenome shotgun sequencing**

Microbial cells in the water samples were collected on 0.2-μm-pore-size cellulose acetate membrane filters (ADVANTEC, Tokyo, Japan) and stored at –80 °C until DNA extraction. Total DNA was extracted from one-eighth sections of the filters as described previously [11] and purified using the UltraClean Microbial DNA Isolation Kit (QIAGEN, USA) according to the manufacturer's instructions. The purified DNA was fragmented using a Covaris S220 ultrasonicator (Covaris, Woburn, MA, USA) to a target fragment size of 500 bp. The shearing conditions were as follows: peak power, 50; duty factor, 20%; cycles per burst, 200; and treatment time, 60 s. Sequencing libraries were prepared from the fragmented DNA using the KAPA EvoPrep Kit (Roche, Basel, Switzerland) according to the manufacturer's instructions, with modifications to the bead purification steps. AMPure XP beads (Beckman Coulter, Brea, CA, USA) were used for bead purification, and the final purification was performed twice at a bead-to-sample ratio of 0.7×. The resulting libraries were sequenced on the Element AVITI platform (Element Biosciences, San Diego, CA, USA) using the AVITI 2 x 300 Cloudbreak Sequencing Kit.

##### **Taxonomic and functional analyses of metagenomic short reads**

Shotgun metagenomic sequencing yielded an average of $22.47 \pm 0.70$ million sequences per sample. Quality filtered metagenomic short-reads generated through the BBDuk utility within BBTools v36.92 (https://sourceforge.net/projects/bbmap/) were classified using SingleM [12] against the Genome Taxonomy Database R232 [13]. Forward reads that passed quality filtering and were $\geq 130$ bp long were screened for functional genes using DIAMOND blastx [14] against a curated and regularly updated database [15] containing 62 metabolic marker genes. These genes capture major pathways involved in energy conservation, carbon fixation, phototrophy, and the cycling of H<sub>2</sub>, CO, methane, sulfur, nitrogen, and iron. Search parameters were optimised for each gene, with a minimum query coverage threshold of 80% and identity thresholds of 80% for *psaA*; 75% for *hbsT*; 70% for *atpA* and *psbA*, *isoA*, *ygfK*, and *aro*; 60% for *amoA*, *mmoA*, *coxL*, genes for [FeFe]-hydrogenases, *nrxA*, *rbcL*, and *nuoF*; and 50% for all remaining genes. To estimate the relative abundance of community members harbouring each gene, read counts were normalised to reads per kilobase million, as previously described [16, 17]. Afterwards, short reads that were functionally classified as *cooS*, *coxL*, or NiFe-hydrogenases were taxonomically classified using Read Annotation Tool [18, 19] against the Genome Taxonomy Database R232 [13].

##### **16S rRNA gene amplicon sequencing and analysis**

The V4–V5 region of the microbial 16S rRNA gene was amplified using the EUB530F–U907R primer pair [20] as described previously [11]. Briefly, the first PCR was performed for 25–35 cycles, followed by treatment with Exonuclease I and shrimp alkaline phosphatase (Affymetrix, Santa Clara, CA, USA) to remove excess dNTPs and oligonucleotide primers. The PCR products were then diluted 100-fold and subjected to a second PCR to attach 8-base barcode sequences. The resulting sequencing libraries were purified twice using AMPure XP beads (Beckman Coulter, Brea, CA, USA) and sequenced on the Element AVITI platform (Element Biosciences, San Diego, CA, USA) using the AVITI 2 × 300 Cloudbreak Sequencing Kit.

Demultiplexed paired-end reads were processed using QIIME 2 release 2026.4 [21]. Primer sequences were removed using q2-cutadapt, and read pairs lacking the expected primers were discarded. The trimmed reads were denoised using the q2-dada2 denoise-paired method [22], with forward and reverse reads truncated at 270 and 230 nt, respectively, based on per-base quality profiles. ASVs were taxonomically classified against the SILVA release 138 full-length SSU rRNA reference database using the q2-feature-classifier classify-consensus-vsearch method [23–25]. Relative abundances from phylum to species level were calculated from the ASV feature table as the proportion of reads assigned to each taxon within each sample.

##### ***Ex situ* seawater incubation**

*Ex situ* CO and H<sub>2</sub> oxidation assays were performed using seawater collected at 5, 300, and 1,000 m from the Challenger and Sirena Deeps. All sample processing, microcosm preparation, and incubations were conducted onboard immediately after CTD recovery. Native seawater was assayed directly, and concentrated cell suspensions were prepared in parallel. Approximately 10 L of seawater was filtered through 0.2-μm, 47-mm polycarbonate membrane filters (ADVANTEC) using an EZ-Stream pump. Retained cells were resuspended in seawater from the corresponding depth to approximately 500 mL, so each concentrate was prepared from a 20-fold larger volume of seawater. Based on measured cell abundances, the cell concentration factors were 2.5–3.9, 3.2–5.2, and 1.6–2.0 for samples collected at 5, 300, and 1,000 m, respectively (Table S5).

Microcosms were prepared by dispensing 85 mL of native or concentrated seawater into 125-mL glass serum vials, leaving an approximately 40-mL ambient-air headspace. Vials were sealed with butyl-rubber stoppers that had been pretreated by boiling in 0.1 N NaOH for 6 h and secured with aluminum crimps as described previously [26]. The headspace was amended with 10 mL of a mixed gas standard containing 10 ppm of gases, resulting in initial headspace mixing ratios of approximately 2 ppm. For each depth and concentration treatment, three live replicates and one autoclaved control were prepared. Sterile controls were autoclaved at 121°C for 20 min, cooled in water, and their headspaces were fully exchanged with ambient air in a biosafety cabinet before gas addition.

Microcosms were incubated horizontally in the dark at 30°C, 15°C, and 4°C for the 5-, 300-, and 1,000-m samples, respectively, with passive agitation from vessel motion. Vials were protected from light using aluminium foil and black plastic bags. For the Challenger Deep assays, the initial headspace sample was collected immediately after gas addition. For the Sirena Deep assays, the microcosms were equilibrated at their respective incubation temperatures for 10 h before initial sampling. At each time point, 2 mL of headspace gas was withdrawn and stored in a 3-mL silicone-sealed vial temporary and analysed within two days. On 22 January 2026 (day 11.3 for Challenger and day 8.0 for Sirena samples), the 5-m microcosms were transferred from 30°C to 25°C because the 30°C incubator became unavailable, whereas the 300- and 1,000-m microcosms remained at 15°C and 4°C, respectively. CO and H<sub>2</sub> were quantified using gas chromatography as described above.

#### Calculation of trace gas oxidation rates and power yields

Trace gas oxidation rates and associated power yields were calculated as described [26], with modifications described below. For each microcosm, the measured headspace mixing ratio of CO or H<sub>2</sub> was converted to the total gas in the vial, defined as the sum of the gas-phase and dissolved fractions. The gas-phase amount was calculated using the ideal gas law, and the dissolved amount was calculated using the Bunsen solubility coefficient for each gas. Solubility coefficients were calculated from the previously established equations [3] using the measured *in situ* salinity and the incubation temperatures of 30, 15, and 4°C for samples collected at 5, 300, and 1,000 m, respectively.

A replicate was classified as exhibiting detectable oxidation when the control-corrected Theil-Sen slope was $< -0.02 \text{ nmol day}^{-1}$ relative to its paired heat-killed control, with classifications additionally confirmed by manual inspection (Table S5). Replicates that did not meet this criterion were classified as non-detects and reported as N.D. and assigned as a value of zero for statistics. For replicates exhibiting detectable oxidation, the active consumption interval was defined manually as the period between the end of any initial lag phase and the onset of the plateau.

Oxidation kinetics were described using a first-order model:

$$C(t) = C_0 \exp(-kt)$$

where $C(t)$ is the total gas at time $t$, $C_0$ is the fitted initial gas, and $k$ is the apparent first-order rate constant. The model was fitted to measurements within the active consumption interval. Volumetric oxidation rates were calculated as:

$$v = kC$$

where $C$ is the *in situ* dissolved concentration of CO or H<sub>2</sub> measured at the corresponding depth and station. When measurements were available from two CTD casts at the same nominal depth, their mean was used for $C$. For concentrated-cell incubations, $k$ was divided by the measured cell concentration factor to express oxidation activity on a native-seawater basis.

Cell-specific oxidation rates were calculated by normalising volumetric rates to the estimated abundance of cells carrying the relevant oxidation genes. Gene-carrying cell abundance was estimated as described above.

The energetic yields of CO and H<sub>2</sub> oxidation were estimated using the thermodynamic framework of Lappan *et al.* (2023). Gibbs free energies under *in situ* conditions were calculated as:

$$\Delta_r G = \Delta_r G^\circ + RT \ln Q_r$$

Standard Gibbs free energies of reaction were $-237 \text{ kJ mol}^{-1}$ for H<sub>2</sub> oxidation ($\text{H}_2 + 0.5 \text{ O}_2 \rightleftharpoons \text{H}_2\text{O}$) and $-257 \text{ kJ mol}^{-1}$ for CO oxidation ($\text{CO} + 0.5 \text{ O}_2 \rightleftharpoons \text{CO}_2$). Reaction quotients were calculated from the *in situ* dissolved concentrations of H<sub>2</sub>, CO, O<sub>2</sub>, and CO<sub>2</sub>. Dissolved CO<sub>2</sub> concentrations were estimated from dissolved inorganic carbon concentrations reported for the Mariana Trench [27] and the *in situ* pH profile [28] using PyCO2SYS [29] at the corresponding *in situ* temperature, salinity, and pressure.

Bulk power yields in $\text{W L}^{-1}$ were calculated as the product of the volumetric oxidation rate in $\text{mol L}^{-1} \text{ s}^{-1}$ and the absolute Gibbs free energy yield in $\text{J mol}^{-1}$. Cell-specific power yields in $\text{W cell}^{-1}$ were calculated from the corresponding cell-specific oxidation rates. A maintenance power requirement of $1.9 \times 10^{-15} \text{ W cell}^{-1}$ was used, corresponding to the median endogenous power requirement of 121 organoheterotrophic bacterial isolates measured at 20°C, as reported and subsequently reanalysed [26, 30]. Differences among depths were tested separately for CO and H<sub>2</sub> and for each oxidation-rate metric. Statistical analysis was performed using the Kruskal-Wallis test and Dunn's post hoc test with Holm-Bonferroni adjustment. All calculations and statistical analyses were performed in Python 3 using SciPy [31] and scikit-posthocs.

#### Supplementary Figures

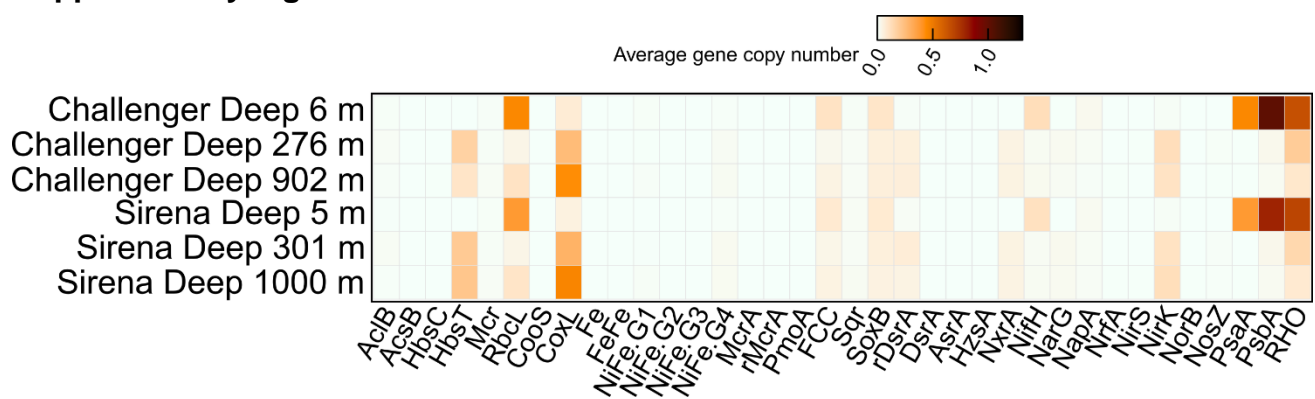

**Fig. S1 Depth-dependent distribution of functional genes in Mariana Trench water-column metagenomes.**

Heatmap showing the estimated average copy numbers of genes associated with phototrophy, carbon fixation, trace-gas oxidation, sulfur and nitrogen cycling, and respiratory energy conservation in metagenomes from Challenger Deep (6, 276, and 902 m) and Sirena Deep (5, 301, and 1,000 m). Rows represent individual metagenomes and columns represent functional marker genes. Color intensity indicates average gene copy number, with darker colors denoting higher values. AcIB, ATP-citrate lyase beta subunit; AcsB, acetyl-CoA synthase; HbsC and HbsT, crenarchaeotal and thaumarchaeotal 4-hydroxybutyryl-CoA synthases, respectively; Mcr, malonyl-CoA reductase; RbcL, large subunit of ribulose-1,5-bisphosphate carboxylase/oxygenase; CooS, anaerobic [NiFe]-carbon monoxide dehydrogenase; CoxL, form I [MoCu]-carbon monoxide dehydrogenase large subunit; Fe, [Fe]-hydrogenase; FeFe, [FeFe]-hydrogenase; NiFe G1-G4, groups 1-4 [NiFe]-hydrogenases; McrA, methyl-coenzyme M reductase alpha subunit; rMcrA, reverse methyl-coenzyme M reductase alpha subunit; PmoA, particulate methane monooxygenase subunit A; FCC, flavocytochrome c sulfide dehydrogenase; Sqr, sulfide:quinone oxidoreductase; SoxB, thiosulfohydrolase; rDsrA, reverse dissimilatory sulfite reductase subunit A; DsrA, dissimilatory sulfite reductase subunit A; AsrA, anaerobic sulfite reductase subunit A; HzsA, hydrazine synthase subunit A; NxrA, nitrite oxidoreductase subunit A; NifH, nitrogenase iron protein; NarG, membrane-bound nitrate reductase alpha subunit; NapA, periplasmic nitrate reductase catalytic subunit; NrfA, ammonifying nitrite reductase; NirS, cytochrome cd1 nitrite reductase; NirK, copper-containing nitrite reductase; NorB, nitric oxide reductase subunit B; NosZ, nitrous oxide reductase; PsaA, photosystem I core protein; PsbA, photosystem II D1 protein; and RHO, energy-converting microbial rhodopsin.

#### A Challenger Deep

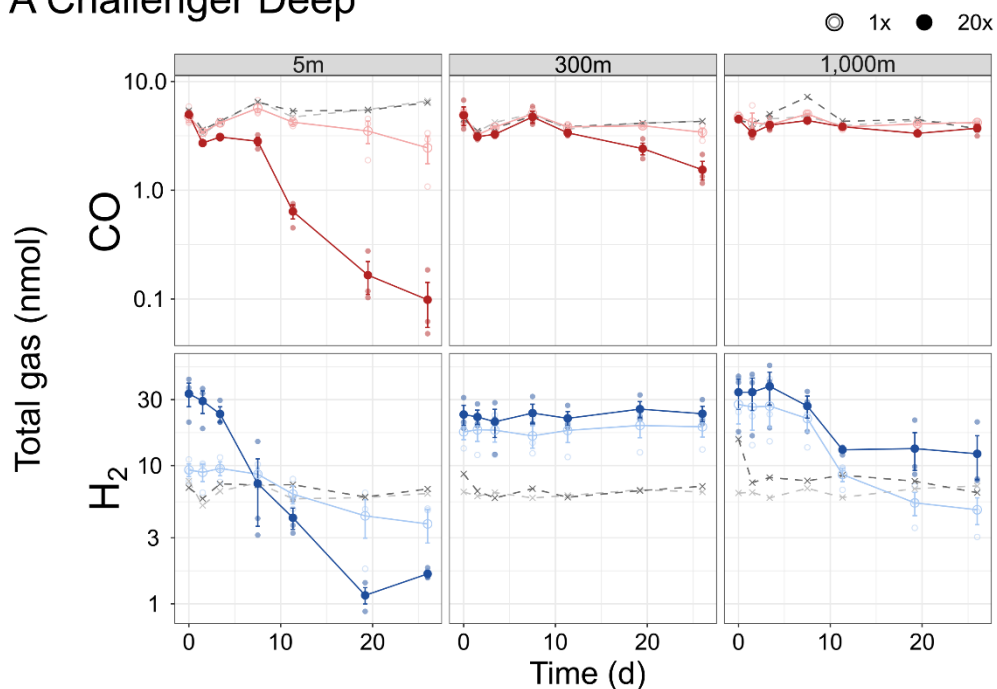

#### B Sirena Deep

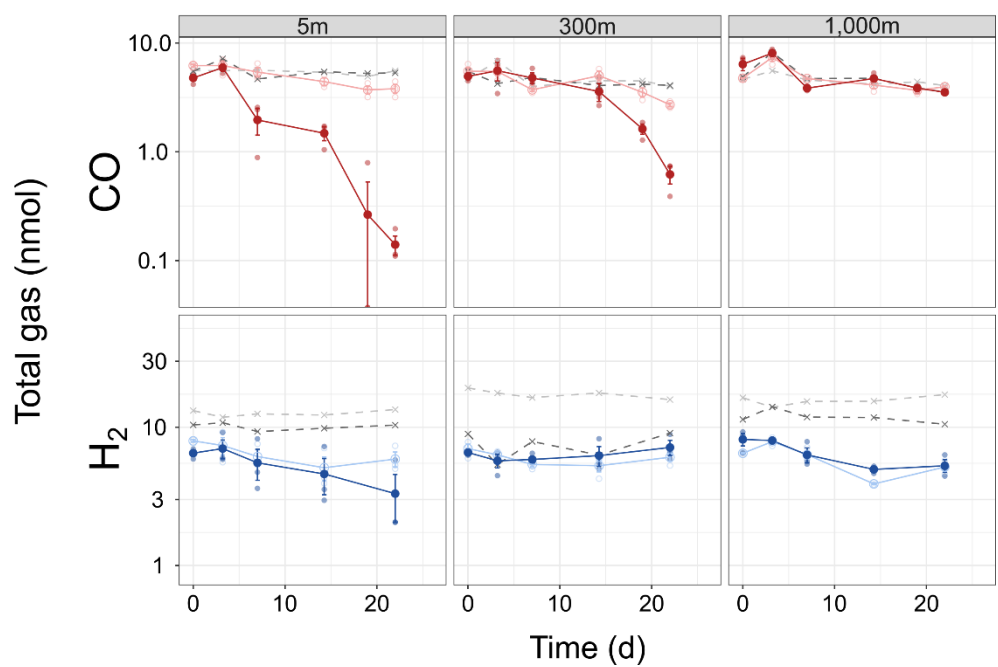

**Fig. S2 Changes in the total amount of CO and H<sub>2</sub> during seawater incubations.** Seawater was collected from 5, 300, and 1,000 m in (A) Challenger Deep and (B) Sirena Deep. CO (red) and H<sub>2</sub> (blue) are shown in the upper and lower rows of each panel, respectively. Total gas represents the sum of the headspace and dissolved fractions and is plotted on logarithmic vertical axes. Open and filled circles indicate unconcentrated seawater (1x) and concentrated-cell treatments prepared from a 20-fold larger seawater volume (20x), respectively. Small symbols represent individual microcosms, larger symbols connected by solid lines represent means, and error bars indicate standard deviations ($n = 3$). Gray crosses connected by dashed lines indicate heat-killed controls. Headspace were amended to approximately 2 ppm CO and H<sub>2</sub>. Microcosms were incubated at 30, 15, or 4°C for seawater collected from 5, 300, or 1,000 m, respectively.
